# Genome-resolved metagenomics reveals cohesive metabolic dynamics, metal resistance genes (MRGs) and biogeochemical cycling in a hot spring system

**DOI:** 10.1101/2022.03.21.485241

**Authors:** Shekhar Nagar, Meghali Bharti, Ram Krishan Negi

## Abstract

The hot spring microbiome is a complex assemblage of micro- and geochemistry of this spring have been and macro- organisms, however, the understanding and projection of enzymatic repertoire that access earth’s integral ecosystem processes remains limited. The composite microbial communities drive global carbon, sulfur, oxygen, iron and nitrogen cycles and other metabolic mechanism involved in heavy metal tolerance and degradation. Interestingly, the Khirganga hot spring microbiome displayed an astounding taxonomical diversity revealed by examination of 41 high and medium qualified metagenome-assembled genomes (MAGs) from at least 12 bacterial and 2 archaeal phyla. Over 1749 genes putatively involved in crucial metabolism of elements *viz*. nitrogen, phosphorous, sulfur and 598 genes encoding enzymes for metals resistance from cadmium, zinc, chromium, arsenic and copper. The MAGs also possess 229 biosynthetic gene clusters dominated by bacteriocins and terpenes could be exploited in medicinal industries. Their metabolic roles found to be altering linkages in biogeochemical cycles and explored a discerned rate of carbon fixation exclusively in archaeal member *Methanospirillum hungatei*. Higher Pfam entropy scores in *Proteobacteria* members highlighting their major contribution in sequestration of ammonia, nitrate and sulfate components as electron acceptors. Through these results, we could postulate that few novel organisms within the community can conduct multiple sequential redox conversions and also considered in reducing emergent difficulties of waste water treatment plants and biotechnological applications.

## Introduction

The hot springs are terrestrial hydrothermal systems harbored with complex *in-situ* chemistry, temperature, pH, and biogeography appeared to be major drivers in forming diverse microbial composition (Skirnisdottir *et al*., 2000, Mathur *et al*., 2007, Pearson *et al*., 2008). High temperature may result in elevated levels of geochemical and physio-chemical deposits. The microbes succeeded to flourish in these conditions by converting chemicals into cellular energy are termed as extremophiles (Cowan *et al*., 2012). Throughout the normal environment, microbes use oxygen for gaseous exchange and derive energy from the earth organic matter, while in extreme thermic environments where O_2_ reduced down to nanomolar levels, only extremophiles dominated or co-evolved utilizing organic and inorganic electron acceptor in several metabolism pathways (Merino *et al*., 2019). A large number of cultivable studies based on 16S rRNA gene analyses have revealed several populations belong to Bacteria and Archaea domains that are phylogenetically distinct from characterized cultures (Yilmaz *et al*., 2016). Since majority of the microbes are uncultivable in hot springs, our recent understanding about the ecophysiology of these uncultivable organisms has become conventional through the omics approaches such as single-cell genomics, de novo metagenomic assembly and genome-resolved metagenomics (Sheik *et al*., 2014, Jochum *et al*., 2018). Success of early genome-resolved metagenomics remained limited due to low sequencing depth of metagenomes (Kunin *et al*., 2008). However, updated high-throughput sequences have refined the sequence quality in low cost and now became applicable to the environmental metagenomes with relatively higher microbial complexities (Yeoh *et al*., 2015). The complex microbial community and functional dynamics in meso-thermic hot spring including genes of metal resistance, abundance of secondary metabolites in Himalayas are poorly understood

In this study, we conducted reconstruction of 41 metagenomes assembled genomes (MAGs) from metagenome datasets of microbial mat, sediment and water of hot spring located at Khirganga, Himachal Pradesh, India. The hot spring is characterized by moderate water temperature (42-60 °C) and geochemistry of this spring have been studied in detail (Nagar *et al*., 2021a). We expanded the bacterial and archaeal diversity performing exchange of inorganic ions such as nitrate, phosphate, sulfate and iron that are hallmarks of many environmental microbes. Also, current study at the community level demonstrated that the inter-organism interactions are the major opinions in the biogeochemical cycling of nitrogen (N), carbon (C), oxygen (O), sulfur (S), iron (Fe) metabolism with hot spring microbiome. We attempt to trace the utility of genomes coupled with metal resistance genes (MRGs) in the biotechnology processes and bioactive secondary compounds of bacteria. The study will readily improve the understanding of composite relationship between bacteria owning MRGs in metal contaminated hot spring and also underlies the exploration of adaptive mechanism of these MAGs in multi-metal contaminated environment.

## Material and methods

### Metagenomic assembly, coverage and binning

The details of sampling and raw data are specified in details in Nagar *et al*. (Nagar *et al*., 2021a). Each individual metagenome was assembled *de novo* using IDBA-UD (Peng *et al*., 2012). The individual assembly across all the 6 samples increased the threshold of reconstruction limiting the strain heterogeneity in replicates. Within each metagenome, the sequence alignment and binary alignment maps were constructed through SAMtools (Li *et al*., 2009), indexing, sorting the reads with Bowtie2 (Langmead & Salzberg 2012) resulted further used to reconstruct bins from assembled contigs. In order to visualize the coverage of assembly, the metagenomic reads were mapped over contigs using Analysis and Visualization Platform for Omics Data (anvi’o) interactive interface (Meren *et al*., 2017), where a blank-profile for sample sequences over contigs >1 kbp was constructed. The microbial genomes were deconvoluted from the metagenomes by combining contigs based on tetra-nucleotide frequency and genome abundance probabilities using Metagenomic Binning with Abundance and Tetranucleotide Frequencies (MetaBAT) v2 (Kang *et al*., 2019) using the following parameters *minContig* (minimum contig size) =2500 bp, and *minS* (minimum score of edge for binning) =60.

### Bin curation, completeness assessment, phylogenetic interference and average nucleotide identity of MAGs

The completeness, heterogeneity and contamination of individual bins were assessed using lineage_wf and quality assessment in CheckM v1.0.18 (Parks *et al*., 2014). These bins were integrated into tree of life along with 3171 microbial genomes based on universal marker protein sequences using PhyloPhlAn v2.0 (Segata *et al*., 2013). Later, the taxonomic placements were done on an open-source platform Genome Taxonomy Database GTDB-Tk (Parks *et al*., 2020, Chaumeil *et al*., 2019) which assign domain to the bins using 120 bacterial and archaeal specific marker genes and further classified on the basis of relative evolutionary divergence (RED) and average nucleotide identity (ANI) with 24,706 microbial genomes present in the database using FastANI (Jain *et al*., 2018) and pplacer v1.1 (Matsen *et al*., 2010). Also, the ANI of each metagenomic assembled genome (MAG) with its nearest neighbor was plotted against the fraction of aligned genome to study the evolutionary divergence of each MAG with its respective neighbor and the trends were compared among all obtained phylum.

### Genomic annotation reveals functional differentiation among the MAGs

The MAGs were annotated using prokka v3.0 (Seemann, 2014) and amino acid sequences were mapped on NCBI COG database using rpsblast v2.2.15 (Altschul *et al*., 1990) to predict functional categories. The COG results were plotted with circular stack bar plot in R (R Development Core team). Next, the MAGs were searched with specific annotations of complementary metabolism included sub classification of nitrogen, phosphorous, sulfur and iron metabolism genes using RASTtk pipeline (Brettin *et al*., 2015) and prodigal (Hyatt *et al*., 2010). The amino acids of bins qualified min. alignment length (150 aa) were searched as references to retrieve metal resistance gene (MRGs) in MAGs using DIAMOND (reference) at e-value < 1 x 10^-10^, identity >60% and coverage >80% against BacMet database (v2.0, experimentally confirmed resistance genes, 2018, Pal *et al*., 2014). Further, the coding sequences of each MAG was searched against biosynthetic gene clusters (BGCs) hidden Markov models using antibiotic and secondary metabolites analysis shell (antiSMASH) v4.2.0 software (Bin *et al*., 2017) which accurately identifies gene clusters encoding secondary metabolites from known in-built annotated domains and modules of broad chemical classes.

### Inference of putative proteins involved in Biogeochemical cycles

Multigenomic Entropy Based Score pipeline (MEBS) described as an integrated mathematical approach framed with relative entropy-based scores (abundance) of essential ecological elements Nitrogen (N), Sulfur (S), Oxygen (O), Carbon (C) and Iron (Fe) in an ecosystem. Also, it is an open-source pipeline benchmarked with dataset of 2107 non-redundant microbial genomes from RefSeq and 935 metagenomes from MG-RAST and used to evaluate, compare and quantify the biogeochemical cycles in omic’s datasets (De Anda *et al*., 2017). The relative entropy-based scores of MAGs provided information on shifts in microbial-mediated biogeochemical cycling and bottom-up effects on entire hot spring. It was computed by using built-in function of MEBS hmmsearch against protein domains of Pfam v3.0 database that were involved in N, S, O, C and Fe cycles with a restrictive false discovery rate (FDR) of 0.001. Further, the normalized Pfam entropy scores, completed and clustered metabolic processes in MAGs were visualized by using mebs_vis.py in MEBS.

## Result and Discussion

### Quantifying contigs coverage for recovering

We used genome-resolved metagenomics approach to evaluate bacterial and archaeal lineages in microbial mat, sediment and surface water from a meso-thermic hot water spring at Khirganga, Parvati Valley, Himachal Pradesh, India (Shirkot and Verma 2015, Nagar *et al*., 2021a). We calculated the sequence composition and differential coverage by mapping the reads over the contigs to avoid misleading conclusion of assembled contigs using anvi’o v2 (Meren *et al*., 2015). Approximately 10.2 million [M] reads (total reads 11[M]) in KgM1, 14.1 (15.2 [M]) in KgM2, 6.6 (14.8 [M]) in KgS1, 6.3 (14.5 [M]) in KgS2, 11.2 (14.4 [M]) in KgW1, 10.9 (14.1 [M]) in KgW2 showed accurate coverage up to 10x where more than 95% of the reads were mapped on the assembled contigs in microbial mat, sediment and water showed with Euclidean distance and ward linkage parameters in Figure 1. A low no. of single-nucleotide variants (SNVs) in all the samples suggested there was a less genomic heterogeneity in all the samples and these contigs could be excluded manually in order to improve binning process. The microbial mat and water samples showed large contig abundance for reconstruction while sediment samples revealed a more complex or heterogenous segments in the assembled contigs.

**Figure 1:**
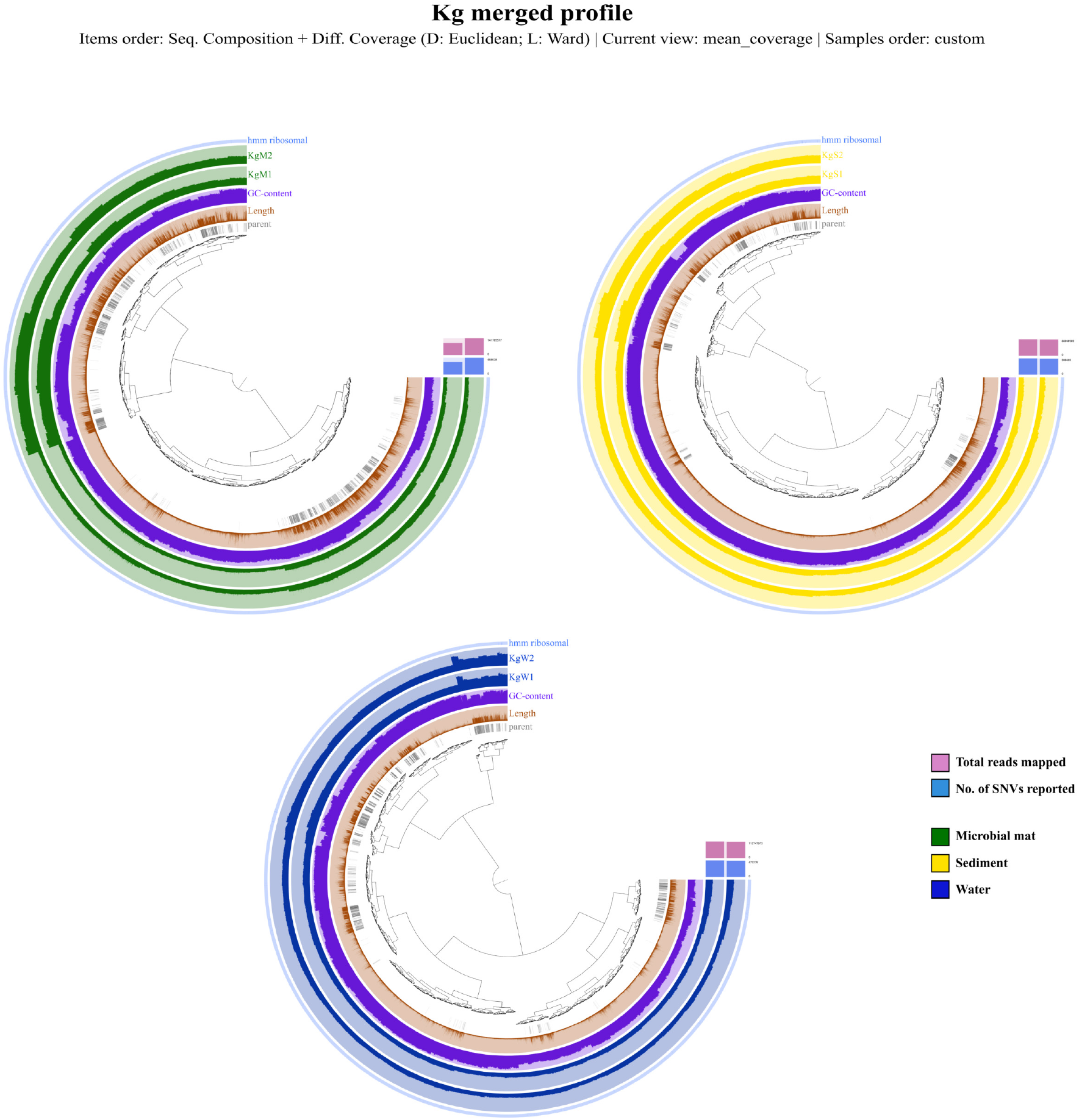
Coverage profiles of metagenomic samples in pairs. Interactive view of the reads and contigs mapped together to represent the accuracy of the assembled contigs formed by using anvi’o metagenomic workflow. The assembled contigs intensity shown in green for microbial mat, yellow for sediment and blue for water samples

### Refining, identifying and classifying Metagenome-Assembled Genomes (MAGs)

Initially, the automatic binning process resulted in 196 bins (s.d. 24.5) in microbial mat, 279 bins (s.d. 2.5) in sediment and 221 bins (s.d. 0.5) in water samples thereafter the bins that were contaminated, heterogenous, incomplete and shorter than 0.5 Mbp were removed manually. Only 41 MAGs were qualified at the threshold set by the Minimum Information about a Metagenome-Assembled Genome (MIMAG) (Bowers *et al*., 2017) in which 11 out of 41 MAGs had > 90% completion and < 3.64% contamination and other 30 were qualified at >50.7 percent completion and < 5.62% contamination showed in Table 1. The bins were widely distributed in microbial tree of life where 39 bins belonged to 12 different bacterial phyla and 2 were classified into *Archaea (Euryarchaeota, Thaumarchaeota)*. Here, 21 bins belonged to phylum *Proteobacteria,* 3 bins each belonged to *Chloroflexi* and *Defferibacteres*, 2 bins each were classified into *Bacteroidetes, Armatimonadetes, Cyanobacteria* and 1 bin each belonged to *Spirochaetes*, *Ignavibacteriae*, *Actinobacteria*, *Nitrospirae*, *Candidatus Hydrogenedentes* and *Verrucomicrobia* (Figure 2A).

**Figure 2:**
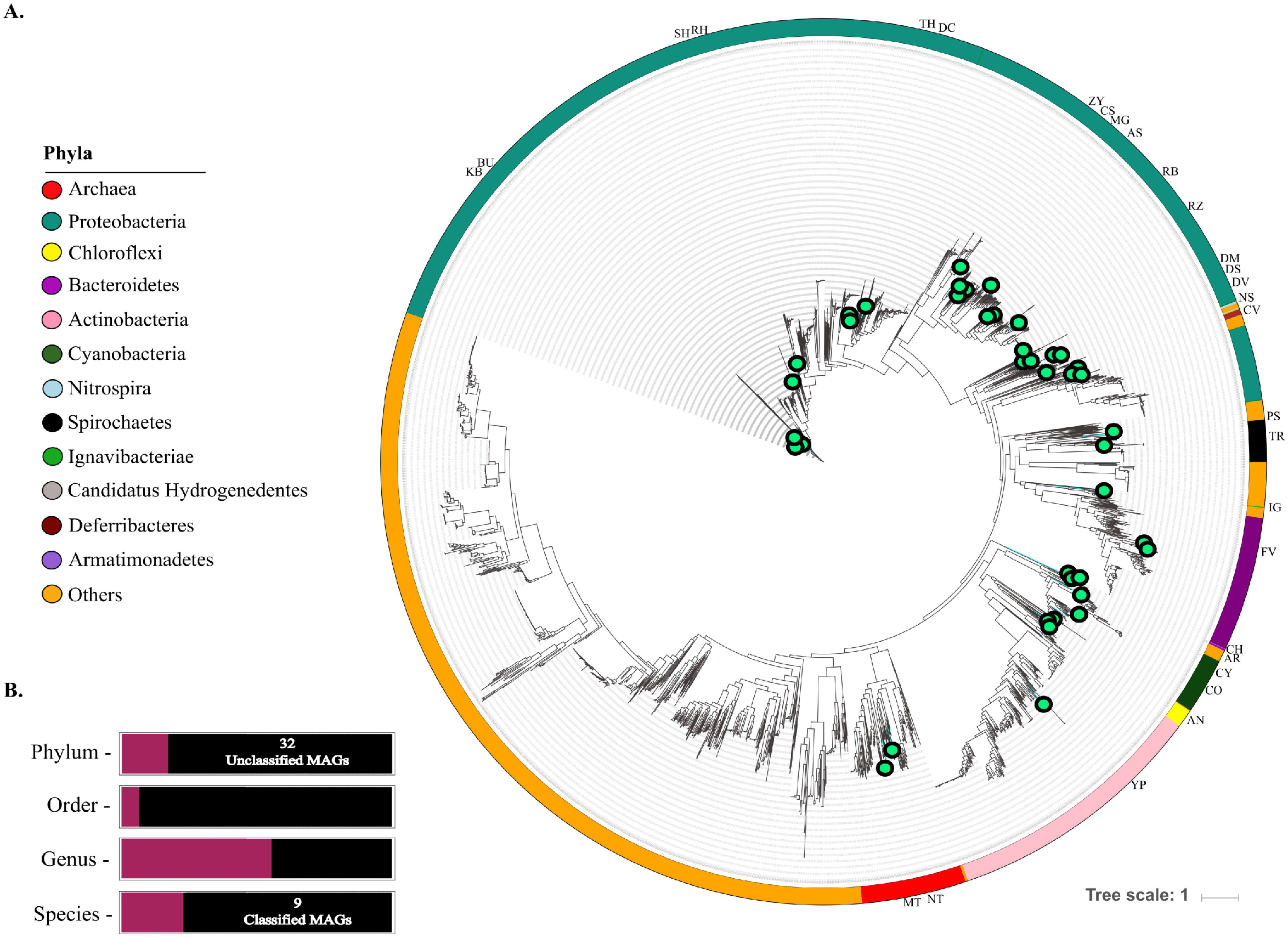
Phylogenetic placement and integration of recovered MAGs in the microbial tree of life having 3,171 bacterial and archaeal genomes using Phylophlan v2. A) The recovered MAGs after quality filter placed as bubbles on the axis adjacent to their neighbors. The outer circle depicting the phylum colors with the genera abbreviations described as *Buttiauxella noackiae* (BU), *Shewanella putrefaciens* (SH), *Candidatus Hydrogenedens* sp. (CH), *Treponema* sp. (TR), *Desulfomonile* sp. (DM), *Anaerolinea* sp. (AN), *Methanospirillum hungatei* (MT)*, Klebsiella quasipneumoniae* (KB), *Coleofasciculus chthonoplastes* (CO)*, Nitrosotenius* (NT), *Ignavibacteria* sp. (IG), *Cyanobacteria bacterium* M5B4 (CY), *Yonghaparkia* sp. (YN), *Prosthecobacter* sp. (PS), *Allorhizobium* sp. (RZ), *Zymomonas* sp. (ZY), *Dechloromonas hydrophilus* (DC), *Asticaccaulis* sp. (AS), *Nitrospira* sp. (NS), *Magnetospirillum* sp. (MG), *Pararheinheimera* sp. (RH), *Thiobacillus* sp. (TH), *Desulfovibrio* sp. (DV), *Caenisprillum bisanense* (CS), *Armatimonadetes bacterium* UBA8783 (AR) *Rhizobiales bacterium* PAR1 (RB), *Desulfarculus* sp. (DS) *Flavobacterium piscis* (FV), *Calditerrivibrio* sp. (CV), *Chloroflexi bacterium* OLB14 (AN). B) The classification of MAGs was estimated through the Genome Database Taxonomy (GTDB-tk). 32 unclassified MAGs were corresponded to those with no associated genome in the database at the species taxonomic level.

**Table 1:**
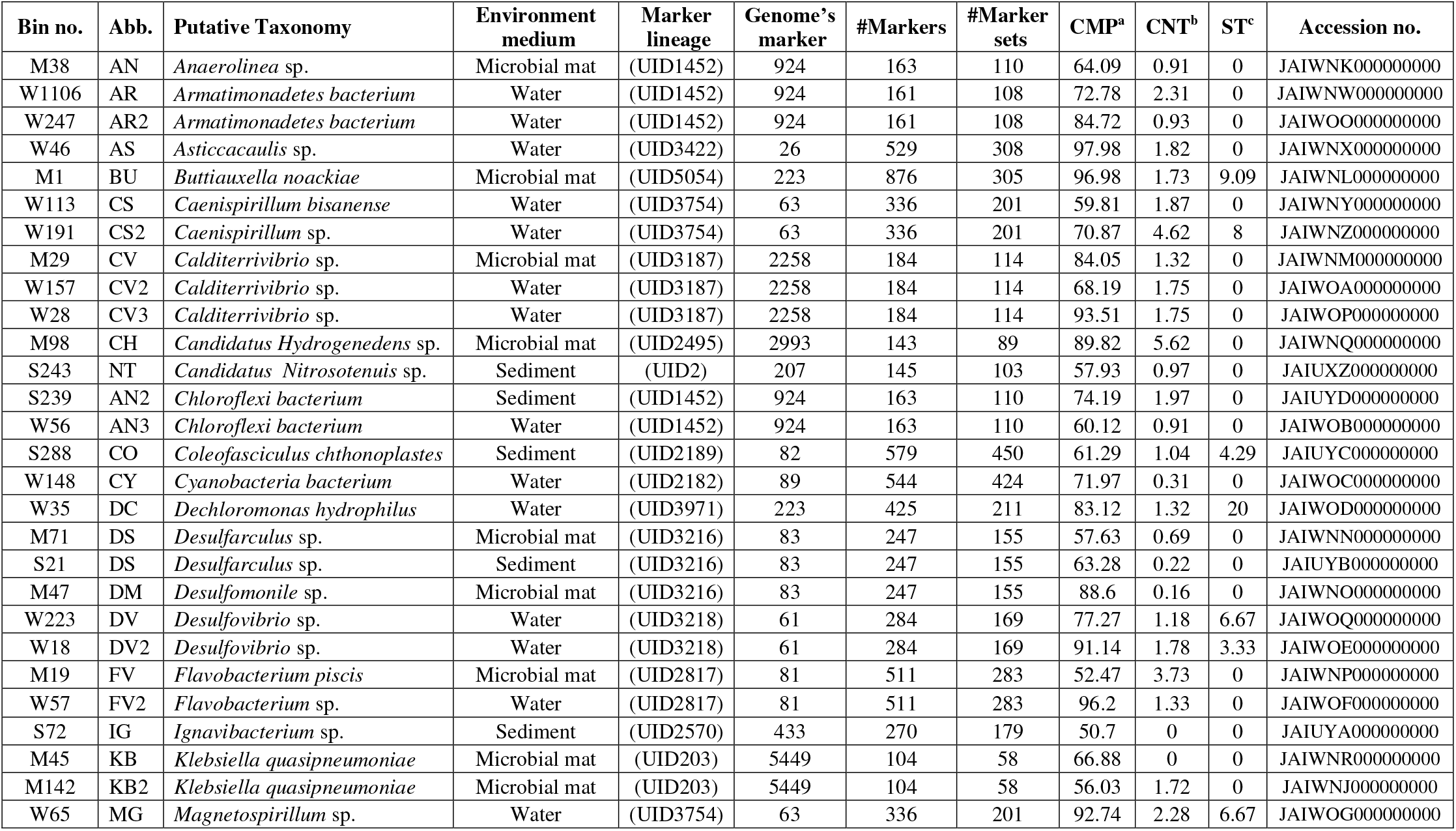
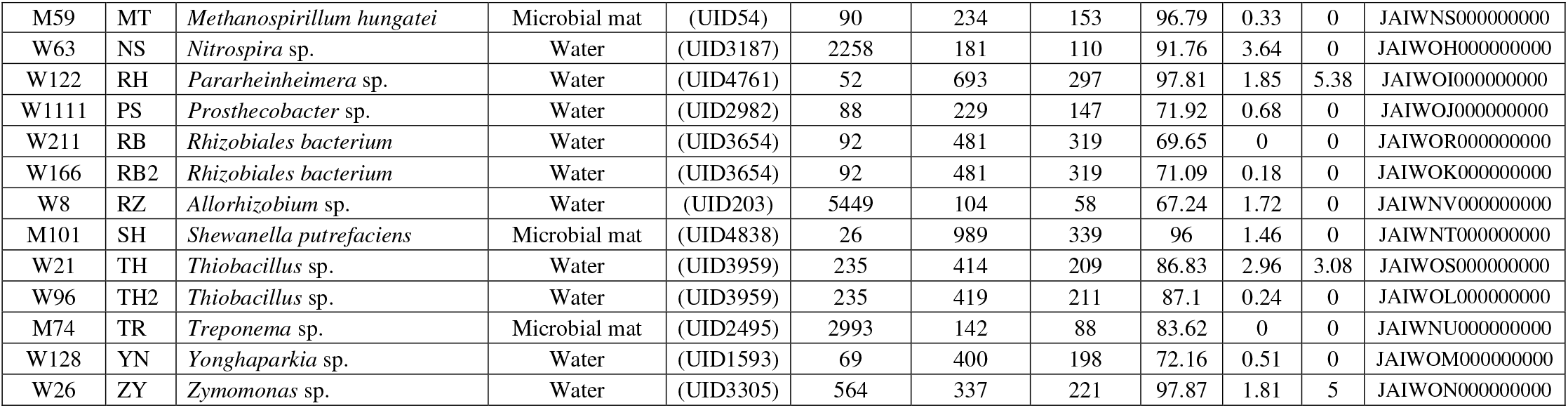
Accessions, and other attributes of Metagenome-assembled Genomes (MAGs)

The protein sequences of the 41 bins were annotated through prodigal using Pfam TIGRfam protein families and align along marker genes to identify the domains. Further, the bins were masked along the concatenated alignment of 23,458 bacterial and 1248 archaeal genomes present in the database therefore, validated scores of ANI, RED and taxonomy placements in the reference tree by using pplacer, we were able to classify 10 individual bins up to species level and others up to genus level validated with scores of the ANI and pplacer placements in the reference tree (Figure 2B; Supplementary material: File S1). The recovered genomes were partitioned distinctively in microbial mat were *Buttiauxella noackiae* (BU), *Shewanella putrefaciens* (SH), *Candidatus Hydrogenedens* sp. (CH), *Treponema* sp. (TR), *Desulfomonile* sp. (DM), *Anaerolinea* sp. (AN), *Methanospirillum hungatei* (MT), two species of *Klebsiella quasipneumoniae* (KB), while in sediment samples *Coleofasciculus chthonoplastes* (CO)*, Nitrosotenius* sp723185 (NT), *Ignavibacteria* sp. (IG) were present and in water *Cyanobacteria bacterium* M5B4 (CY), *Yonghaparkia* sp. (YN), *Prosthecobacter* sp. (PS), *Allorhizobium* sp. (RZ), *Zymomonas* sp. (ZY), *Dechloromonas hydrophilus* (DC), *Asticaccaulis* sp. (AS), *Nitrospira* sp. (NS), *Magnetospirillum* sp. (MG), *Pararheinheimera* sp. (RH), two species of each *Thiobacillus* sp. (TH), *Desulfovibrio* sp. (DV), *Caenisprillum bisanense* (CS), *Armatimonadetes bacterium* UBA8783 (AR) and *Rhizobiales bacterium* PAR1 (RB) were found. Distinctly, the genus *Desulfarculus* sp. (DS) resides in both microbial mat and sediment samples, whereas *Flavobacterium piscis* (FV) and *Calditerrivibrio* sp. (CV) were recovered from both microbial mat and water samples and the only genome reconstructed in sediment and water samples was *Chloroflexi bacterium* OLB14 (AN). The MAGs distributed in all three habitats (Microbial mat n= 12, Sediment n= 5, Water n= 24) were submitted to NCBI shown in Table 1. The analysis provides community-wide insights within microbial mat, sediment and water habitats of hot spring. Proteobacteria, in metagenomic datasets were overrepresented. As a part of this study, we recovered three genomes from *Chloroflexi* bacterium (Id 233189), *Calditerrivibrio* sp. (Id 545685), one from *Coleofasciculus chthonoplastes* (Id 669368), *Zymomonas* (Id 541), *Dechloromonas hydrophilus* (Id 73029) with very few species cultivated and characterized (Wilkins *et al*., 2019, McDaniel *et al*., 2021).

Further, MAGs were plotted with ANI values on x-axis and alignment fraction on Y-axis showed that 18 MAGs in order of RB, SH, CY, BU, AR, FV, AR2, KB, DC, FV2, CS, RZ, AS, RB, CS, ZY and MT were having more than 92% ANI values with their neighbors when searched against 24,706 bacterial and archaeal genomes. While minimum ANI values 72.29 ± 0.20 were calculated in 5 MAGs namely *Desulfomonile* sp. (DM), two *Chloroflexi bacterium* OLB14 (AN), *Anaerolinea* sp. (AN) and *Coleofasciculus chthonoplastes* (CO) (Figure 3A). The genomes with low ANI values were rare and had very less information available. The completeness lied within a range of 60.1-88.6%, they can be categorized into novel MAGs that belonged to a particular genera or species (Supplementary material: File S1).

**Figure 3:**
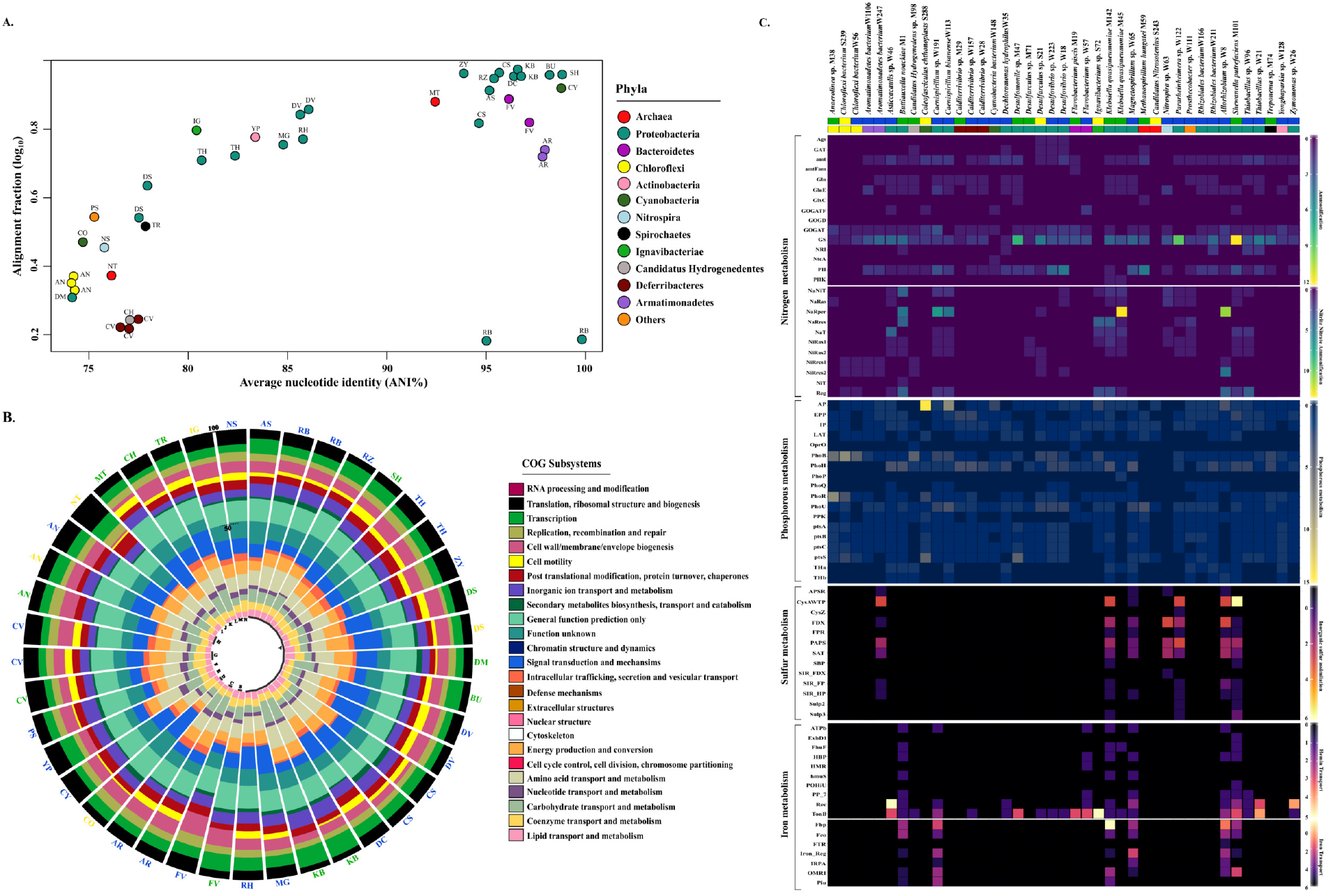
Functional annotations of reconstructed genomes. A) Average nucleotide identity (ANI) values (*x*-axis) plotted against log_10_-transformed alignment length (*y*-axis). Each color-coded dot indicates the best hit between MAG and its closest relative according to the database implemented in GTDB-Tk. The color and abbreviations are denoted to phyla and genera, respectively. B) The circular stack bar plot displaying the functional annotation of the MAGs where each stack is color-coded according to the subsystem percentage determined using best hits with NCBI COG database. C) Heatmap representing the distribution and presence of genes involved in nitrogen, phosphorous, sulfur and iron metabolism, added colored strip with green (microbial mat), yellow (sediment), blue (water) showing the habitat of the genomes reconstructed.

### Genome-specific metabolic reconstructions

The functional profiles of MAGs were annotated by using Prokka v1.12 and the amino acid sequences were blast against the COG protein database using blastp. A circular stack bar plot was constructed for functional differentiation using ggplot2 (Wickham, 2009) in R (R Development Core team). The total number of genes present in MAGs were 86,904 estimated by using Roary (Page *et al*., 2015). The highest number of CDS (coding sequences) were present in *Coleofasciculus chthonoplastes* followed by *Klebsiella quasipneumoniae* and *Buttiauxella noackiae* (Nagar *et al*., 2021b). We observed that the functional COG subsystems were evenly distributed in all the MAGs where mean percentage of 12.42 ± 1.32 of gene sequences encoded to general function prediction only (R), 9.36 ± 3.99 of sequences account for signal transduction mechanisms (T), 9.12 ± 1.91 of sequences annotated to be as amino acid transport and metabolism (E), 8.09 ± 1.77 belonged to function unknown (S) and other COG classes encoded by 8% of sequences (Figure 3B; Supplementary material S1). Further, the annotated amino acids were searched against the specific databases of elements sulfur, phosphate, nitrogen and iron.

#### A) Nitrogen metabolism

In general, element like nitrogen (N) is abundant in atmosphere precipitated in the form of oxides and ammonia salts. Analysis of nitrogen metabolic potential across MAGs revealed a higher abundance of genes involved in nitrite and nitrate reduction and ammonia assimilation. Reportedly, the anerobic respiration by chemoorganoheterotrophic bacteria utilizing nitrate (NO3^-^) as electron acceptor producing ammonia ions (NH4^+^) termed as dissimilatory nitrate reduction to ammonia (DNRA) and this process is prevalent in hot springs (Dodsworth *et al*., 2011, Tripathy *et al*., 2016, Alcamán-Arias *et al*., 2018). So, we investigated the genes involved in dissimilatory reduction of nitrate to ammonia were (n= 181) *NaNiT, NaRas, NaRper, NaRres, NaT, NiRas1, NiRas2, NiRres1, NiRres2, NiT, Reg* which were found to be highly enriched in order by BU> MG> SH> KB> CS (Figure 2C). The denitrification and DNRA are important to equilibrate NO3^-^ in environment nitrogen cycle and provide soluble NH_4_^+^ (Lam *et al*., 2011) in favorable conditions (less NO_3_^-^ than organic carbon), DNRA may achieve higher growth curves on account of efficiently conserving its enzymatic reaction and outcompete denitrifiers (Kraft *et al*., 2011). Here, the habitats of hot spring with low NO3^-^ content revealing the prevalence of proteobacteria involved in active DNRA pathway (Mohan *et al*., 2004, Giacomucci *et al*., 2012). Further, the microbial ammonia assimilation was evaluated as it is essential to assimilate ammonia into organic compounds in maintaining N cycle. The genes involved in ammonia assimilation (n=337) process namely *Ags, GAT, amtFam, Gln, GlnE, GlxC, GOGATF, GOGD, GOGAT, GS, NRI, NtcA, PII, pIIK* (Supplementary material S3; NH_4_^+^ → amino acids, proteins, nucleic acids) were widely distributed in all MAGs in order SH> DM> TH> CS> RH (Figure 3C). The key enzymes *GOGAT* (Glutamate synthase [NADPH]) and *GS* (Glutamine synthase) were present in all the metagenome-assembled genomes except in *Ignavibacteria* and *Candidatus Nitrosotenius* (Figure 3C). The ammonia ions fixed directly into glutamate through alpha-ketoglutarate (NADPH-dependent) reaction by glutamate dehydrogenase (GDH) and indirectly through glutamine synthetase (GS) to glutamate-oxoglutarate amidotransferase (GOGAT) and further synthesized into polyamines, nucleic acids and organic nitrogen (Nagatani *et al*., 1971). The denitrification (n=65) and nitrosative stress (n= 67) pathways were also widely distributed in MG, CS, FV, RZ (Supplementary material: S2). The global chemical N cycle is conserved in the ecosystem and its metabolically active genes monitoring could be pragmatic in detecting baseline nutrient levels and drifts, preventing eutrophication, minimizing noxious effects of ammonia or nitrite and maximizing soil productivity (Xu and Zhou 2004, Jacoby *et al*., 2020).

#### B) Phosphorous metabolism

Phosphorous (P) exists on earth in the form of inorganic phosphate (P_i_), phosphate-containing minerals, organic phosphate esters and their derivatives. In anaerobic conditions, the bacteria that could uptake the phosphorous to synthesis poly-β-hydroxybutyrate called as Phosphorous accumulated bacteria (PAB). The major genes involved in phosphate metabolism (n=807) *AP, EPP, IP, LAT, OprO, PhoHBPQRU, PPK, ptsA, ptsB, ptsC, ptsS, THa, THb* were distributed in all the assembled genomes with maximum no. of copies present in CO, CS, AN, MG (Figure 3C, Supplementary material S3). In limited Pi conditions, alkaline phosphatase (AP) encoded by *PhoA* hydrolyzes the phosphonates releasing Pi in periplasmic space subsequently transported across the inner membrane and assimilated by *PhoBR* genes. The protein synthesis of these genes increased up to 1500 times suggested their important role in scavenging P_i_ from environment during P_i_ starvation conditions, (Willsky & Malamy, 1976, Lu *et al*., 2019). Here, it could be confirmed in all MAGs that if there is limited availability of P_i_ then whole Pi starvation inducible gene expression (Psi) was controlled by the *Pho* regulon utilising IP of all phosphorous sources through Pi-specific transport (*PtsSCAB*) system (Wanner 1994, White & Metcalf, 2007, Zheng *et al*., 2019). Similarly, the other important alkylphosphonate genes (n= 92) *PhnAB, PhnFGHIJKLMNO, resF* were abundant in BU, KB, RZ, MG, CS, FV, RB respectively (Supplementary material S2). The analysis revealed that the genomes (PABs) are utilizing inorganic phosphite, organophosphates and phosphonates under phosphate ions (Pi) starvation conditions (White and Metcalf, 2004). The mode of phosphorous metabolism in all the MAGs except *Nitrospira* sp. was accomplished through degradation of phosphonates and alkyl phosphates. The PABs recovered from the hot spring could be encouraged for removal of total organic carbon and ammonia in well-established Enhanced biological phosphorus removal (EBPR) processes of wastewater treatment plants (White & Metcalf, 2007, Hirota *et al*., 2010).

#### C) Sulfur metabolism

Sulfur is the essential biomolecule and vital to the organism which usually exists in organic forms. Earlier, we revealed the contamination of sulfate ions and dominance of obligatory anaerobic bacterial and archaeal lineages sulfur related microorganisms (SRM) and hypothesized that dissimilatory reduction pathway was favorable in Khirganga hot spring (Nagar *et al*., 2021a). We identified the sulfur metabolism genes present in MAGs were critically important. The MAGs were explored to gain insights into inorganic sulfur assimilation genes (n=110) *APSR, sat, PAPS, SIR_FDX, SIR_FP, SIR_HP, cysAWTP* (transporter) discussed in Supplementary material S3, were abundant in SH, RH, RZ, KB, NS, AS, MG (Figure 3C, Supplementary material S2) responsible for biosynthesis of several amino acids and other essential compounds like vitamins (thiamin, biotin), prosthetic groups (Fe-S clusters). Apparently, we were able to recover sulfate reducing bacteria (SRB) from all three habitats namely *Desulfovibrio* sp. (water), *Desulfarculus* sp. (microbial mat and sediment), *Desulfomonile* sp. (microbial mat) and sulfur oxidizing bacteria (SOB) *Thiobacillus* sp. (water) and we were able to dig out sulfite reductase complex cluster genes (n=90) *DsrA, DsrB, DsrMKJOP* (Supplementary material: S2) in others species such as *Buttiauxella noackiae*, *Shewanella putrefaciens* and *Magnetospirillum* sp. (Hedlund *et al*., 1997, Anantharaman *et al*., 2018). Also, dissimilatory sulfate reduction proteins were found to be present in TH, MG, DM, DV, DS, KB, SH, BU, RB in all three habitats of the hot spring (Supplementary material S3). It is evident that the dissimilatory reduction of sulfate is less complex than assimilatory sulfate reduction and could be attained easily in various sulfate reducing bacteria (SRB) which pass a toxic product sulfide in exchange with water and oxidized to form sulfite and sulfates (Kushkevych *et al*., 2020). The bioengineered SRBs are better candidates to be employed to enhance the sulfate and toxic metals elimination through dissolution and precipitation in acid mine tailings waste (AMTW) treatment plants (Zhang and Wang 2016, Dunbar 2017).

### Abundance of metal resistance genes (MRGs)

To assess the heavy metal contamination in the hot spring reconstructed candidates, we identified the Chromium (Cr), Cobalt (Co), Zinc (Zn), Cadmium (Cd), Copper (Cu) and Arsenic (As) resistance genes in metagenome-assembled genomes. Here, 50% of the genomes in which KB, DS and FV, RH, MG could be represented as key microorganisms contributing in Cr and Co-Zn-Cd (CzcCAB efflux system) resistance (n=321), respectively (Supplementary material S4; Figure 4A) and the genes with their specific functions were described. Resistance to arsenic was mediated by ars operon where structural and catalysis genes *arsABC* (n=100) and regulatory proteins ArsR (repressor) and ArsD (co-regulator of arsRDABC) were present (Supplementary material S4, Wu and Rosen, 1993, Bhattacherjee *et al*., 2000). The genes *arsABC* were enriched in 22 MAGs where maximum was contributed in order by MG, CS, SH, FV (Figure 4A). The regulation of *copCBA* system (n=181) for copper resistance is mediated with expression of CopZ (Chaperon) and CopR (repressor) protein. Here, *copCBA* system was not active but CopZ found to be enriched in order CO, AN, BU, DM, DS, MG, RH and YN only transporters CIA, ClfA and gene HL were detected (n= 36, 12, 22, respectively, Figure 4A). Presence of copper transporting genes (n=96) in MAGs ensuring the homeostasis of Cu (+) proteins and also revealing the appropriate uptake, transport and trafficking of copper as a redox co-factor in catalytic center of enzymes (Kim *et al*., 2008, Adriene *et al*., 2008)). The toxicity of copper in hot spring habitats is ascertain and might be mediated through various other mechanisms in environment (Srivastava *et al*., 2020). The efflux system is preferred to maintain Cu intake and uptake, as an essential process in bacteria (Outten *et al*., 2001; Franke *et al*., 2003). The element copper serves as an essential co-factor for several oxidative stress related enzymes including cytochrome c oxidases, monoamine oxidase, superoxide dismutase, catalases, dopamine-monooxygenase and peroxidases (ATSDR 2002, Harvey *et al*., 2008, Stern 2010).

**Figure 4:**
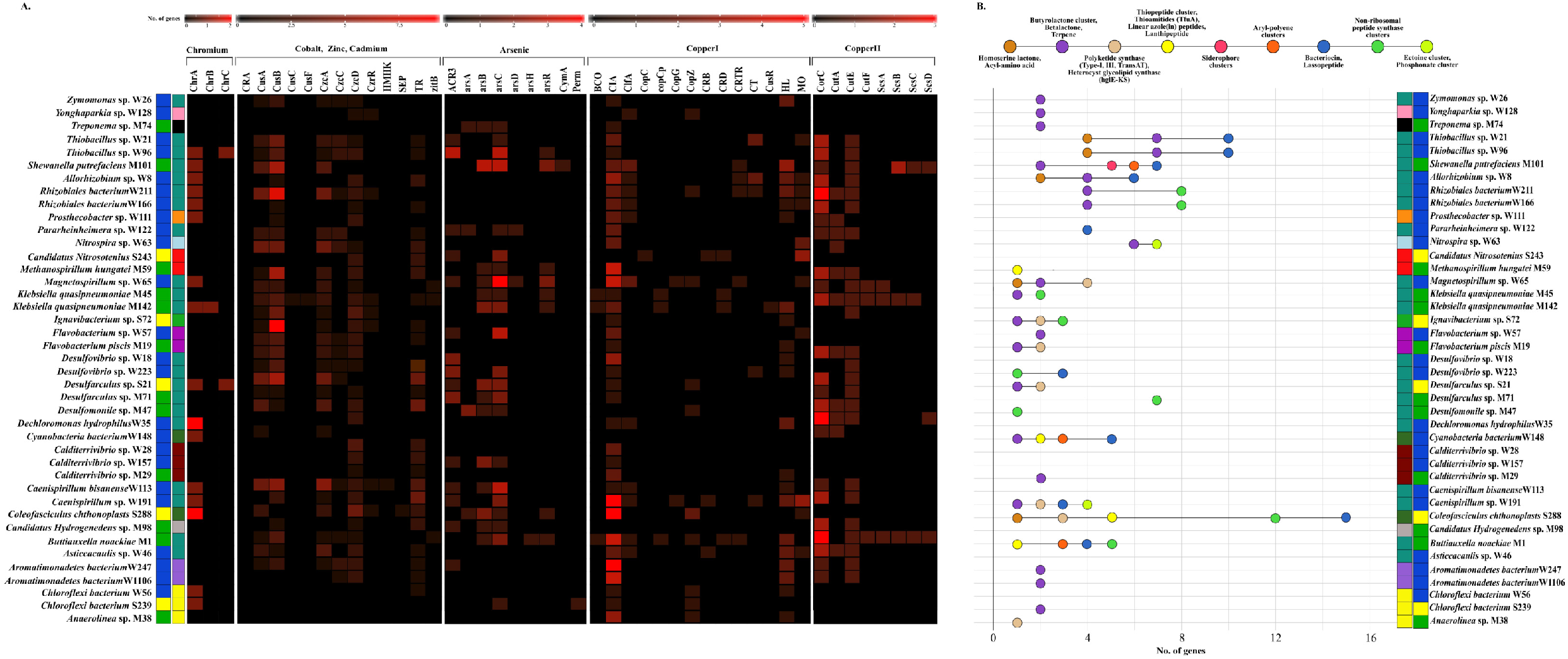
Metal resistance and secondary metabolites estimation of each MAG. A) Heavy metal resistance genes, B) The nodes of lollipop plot here depicting the quantitative and qualitative diversity of secondary metabolites contained by each metagenome-assembled genome reconstructed from the hot spring. Alignment of the genomes with font color green (microbial mat), yellow (sediment) and blue (water) on y-axis and copy number of secondary metabolites biosynthetic gene clusters (BGCs) were aligned on x-axis detected for each MAG using antiSMASH v4.2 software.

Heavy metals are bio accumulative in nature, toxic and persisting in natural setting due to weathering of metal bearing rocks, volcanic eruptions and various agricultural and industrial mining. The microbiota acquire resistance with cumulative metal concentration thresholds irrespective of their essentiality or non-essentiality to the environment. Here, the metal speciation suggested that 75% of the reconstructed MAGs were showing resistance to Cr, Co, Zn, Cd, As, Cu and have evolved to tolerate heavy metal concentrations using various processes includes biosorption, intra/extracellular entrapment and transportation across cell membrane which form the basis of different bioremediation strategies (Ahemad, 2012, Nanda *et al*., 2019, Lin *et al*., 2021).

### Distribution of Secondary metabolites in metagenome-assembled genomes

The Biosynthetic Gene Clusters (BGCs) group of collated genes function together as a molecule produce essential bioactive compounds known as secondary metabolites. Here, we determined 9 secondary metabolites biosynthetic regions (total regions, n=229) in 30 MAGs varying from 0.01 kb to 87.5kb in which (i) bacteriocins, lassopeptide (n= 67) were abundant in *Coleofasciculus chthonoplastes* with other 8 members, (ii) the second highest were butyrolactone cluster, betalactone, terpene (n= 57) enriched in *Coleofasciculus chthonoplastes* with other 17 MAGs; (iii) others encoded for non-ribosomal peptide synthase clusters (NRPS, n=47); homoserine lactone, acyl-amino acid (copy number, n= 12); thiopeptide cluster, thioamitides (TfuA), linear azole(in) peptides, lanthipeptide (n= 9); siderophore cluster (n=1); aryl-polyene clusters (n=11); ectoine cluster, phosphonate cluster in NS and CS2 (n= 11); polyketide synthase (Type-I, III, TransAT), heterocyst glycolipid synthase (hglE-KS) (n= 14) visualized in Figure 4B. The result here showed that 74.6% of the BGC were classified into NRPS-PKS, bacteriocin and terpenes clusters and higher permutations of genome-by-BGCs domains were present in sediment candidates (8.8 _ 19.21%, Wilcoxon’s paired test, p-value < 0.05) than in water (5.6 _ 58.9%) and microbial mat (4.16 _ 21%) genomes. Here, the increase in diversity in the hot spring corresponded to increase in BGCs clusters in microbial communities and the abundance could be hypothesized due to variation in ecological forces such as pH, temperature and bedrock lithology of the hot spring as compared to others (Li *et al*., 2014, Micallef *et al*., 2015). These BGCs are diverse array of bioactive compounds includes growth hormones, antibiotics, quorum sensing, immunosuppressants principally produced by the activation of ambiguous gene clusters which are secreted under extreme conditions (Hibbing et al, 2010). The chemical properties of secreted BGCs were manifesting immunity, controlling overcrowding (bacteriocins), quorum sensing (butyrolactone cluster, betalactone, acyl-amino acid, homoserine lactone), antibacterial, insecticidal, iron acquisition (terpenes, non-ribosomal peptide synthase clusters, polyketide synthase, heterocyst glycolipid synthase, lanthipeptides), safeguarding enzymatic activities (ectoine and phosphonate cluster) in reconstructed genomes (Riddley *et al*., 2008, Song *et al*., 2011, Mark, 2018, Van thouc *et al*., 2019, Caetano *et al*., 2020, Amin & Kannaujiya, 2021). The chemical diversity and gene ontology of these MAGs could be exploited in medicinal industries to improve animal and human health (Ruiz *et al*., 2010, Petit *et al*., 2011, O’Brien and Wright, 2011, Xu *et al*., 2019).

### Significance of MAGs in biogeochemical cycles

The relative abundance of entire biogeochemical cycles (one of the essential integral ecosystem processes) was inferred in all recovered MAGs using MEBS relative entropy scores (H’). The biochemical pathways of S, C, O, Fe and N involved in ecological roles were evaluated based on amino acid sequences for each genome. Domain hits with genomes measured with H’ values close to or greater than 1 correspond to the most informative Pfam domains, whereas low H’ values (zero) indicate non-informative ones. The analysis revealed that the highest MEBS score corresponded to the iron cycle maximum 0.95 in SH lowered down with MG, CS, AN, DM (mean 0.44) revealed the presence of iron oxidation and reduction mechanisms and it also interacts significantly with phosphorous, nitrogen and sulfur cycle (Figure 5C). The second highest entropy scores corresponded to carbon cycle found maximum 0.9 in MT decreased in an order to nearly zero by YN, DS, FV (mean 0.08) suggesting the presence of methyl-coenzyme reductase complex (mcr) and their ability to degrade methane methyl compounds, such as methylamine, a common compound in the hot spring ecosystem (Hua *et al*., 2019). The third highest scores estimated in nitrogen cycle completed in MG (0.83) reducing in order by CS, CO, RZ, NS (mean 0.48) indicated that these MAGs could use nitrogen compounds, such as ammonia or nitrates, as energy sources. The oxygen cycle in MAGs was also being controlled significantly by CO with maximum score of 0.79 followed by MG, CS, CY (mean 0.34) explaining their aerobic role in environment. Lastly, sulfur cycle estimated maximum 0.72 in TH and more than 0.5 in all *δ-proteobacteria* group DV, DM, DS, YN (mean 0.25) indicated the complete pathways of sulfur cycle (Supplementary S5).

**Figure 5:**
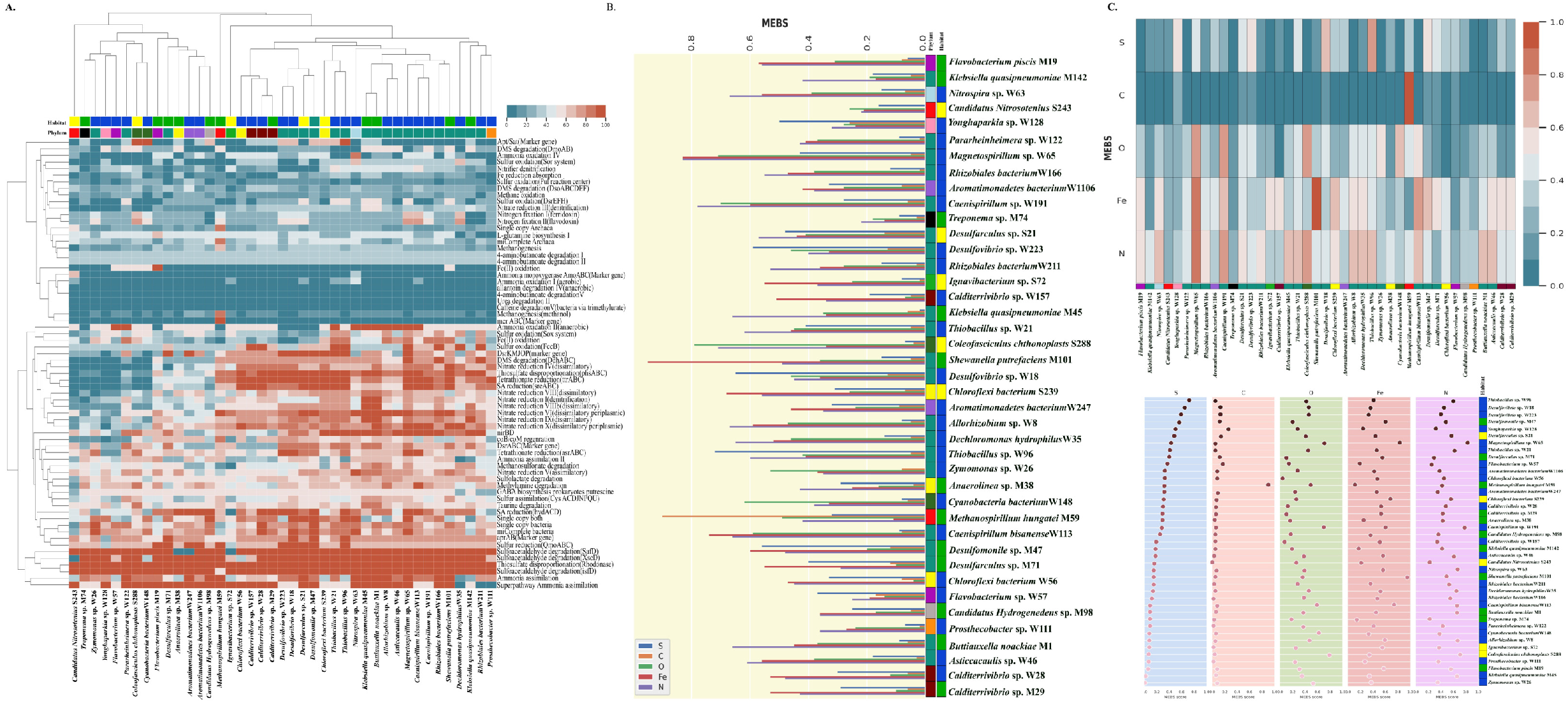
Biogeochemical cycling within the 41 MAGs reconstructed from microbial mat (green), sediment (yellow) and water (blue). (A) Metabolic completeness of essential biogeochemical pathways shown with color gradient, the completed pathways shown in red color and minimal reconstructed in blue color. The strip colored according to the classified phylum of MAGs discussed earlier in tree of life, (B) Comparative depiction of distribution of biogeochemical cycling in each individual MAG, (C) The entropy scores (H’) of N, S, C, O and Fe were estimated across all MAGs and positioning of MAGs with a lower to higher sulfur entropies in order to represent a relative biogeochemical cycling of microbes in different habitat.

Here, the completeness of specific and essential biogeochemical metabolic pathways in the metagenome-assembled genomes, showed by generating a hierarchical clustered heatmap where three pathways thiosulfate disproportionation (rhodonase); sulfoacetaldehyde degradation (isfD); sulfoacetaldehyde degradation (XscD) with 100% completion were present in all the MAGs (Figure 5A, Supplementary material S5). The presence of rhodonase enzyme suggested their role in following substitution transfer pathway resulted in detoxification of hazardous cyanides formed in the hot spring (Gupta *et al*., 2010). The enzyme sulfoactealdehyde belong to short-chain dehydrogenase/reductase (SDR) family was highly abundant which participates in metabolism of taurine and hypotaurine that produces hydrogen sulfide as byproduct (Weinitschke 2010, Rohwerder, 2020). We identified the complete coenzyme M reductase (mcrABC) complex in genome *Methanospirillum hungatei* (MT) which is a key marker enzyme for methane metabolism and involves in degradation of methyl compounds as a source of energy (Hua *et al*., 2019, Shaw *et al*., 2020). The unique metabolism of primitive enzyme ammonia monooxygenase AmoABC was only completed in *Candidatus Nitrosotenius* (NT) and *Nitrospira* sp. (NS) which is involved in producing energy by aerobic oxidation of ammonia to form hydroxylamine and nitrite as a byproduct (Norton *et al*., 2002, Norton *et al*., 2008). The analysis of estimation of biogeochemical cycles gained complete insights in the complex metabolic chemical reactions of nitrogen, sulfur, iron and oxygen cycles in MAGs specified despite their prodigious capacity of molecular transformation. We demonstrated that phylum Proteobacteria candidates have attained maximum biochemical sequestration of ammonia, nitrate and sulfate components as electron acceptors (Figure 5A). The resulted low scores of carbon cycle in all MAGs (except MT) evidencing the minimal degradation of organic compounds by microbial driven processes in the hot spring and surrounding atmosphere (Figure 5C).

## Conclusions

The overall functioning of microbial communities from hot spring has remained indefinite as most of these occur at low abundance or difficult to characterize genomically. We implied genome-resolved metagenomics approach to deconvolute 41 genomes from microbial mat, sediments and water of hot spring. Few organisms such as *Chloroflexi* bacterium (Id 233189), *Calditerrivibrio* sp., *Coleofasciculus chthonoplastes*, *Zymomonas*, *Dechloromonas hydrophilus* are very rare and extracted from habitats of hot spring. We studied and gained insights into metabolic mechanisms of nitrogen, phosphorous and sulfur metabolism in taxonomically diverse metagenome-assembled genomes. The inter-conversion of nitrogen to ammonia to nitrate may be a prominent exchange but through dissimilatory process termed as DNRA. The sulfur metabolism may be larger with complete oxidation and reduction processes. The genomic repertoire suggested the ongoing specific adaptations (MRGs; Czc efflux system, copABC, arsABC) to cope up with extreme standards of chromium, cadmium, zinc, cobalt, arsenic and copper metal resistance. The genomes were also leverage the secondary immunity components (BGCs) mainly bacteriocins, terpenes and NRPS. Further, the study was designed to investigate the alternating roles of microbial taxa in biogeochemical cycles. Here, the metabolic versatility indicate that *Proteobacteria* may occupy essential roles in anaerobic nitrogen, sulfur and iron cycle while *Euryarchaeota* are capable of using simple organic compounds plays part in carbon cycle. The tremendous metabolic individualities and novel phylogenetic diversity in the genomes reported here could be represented as trait-based geothermal models required to predict the effects of changing extreme conditions on biogeochemical cycles. The organisms with complete sulfur, nitrogen, phosphorous metabolism, heavy metal tolerance and BGCs could become potential candidates to be implied in biotechnological and bioremediation applications.

## Availability of Data

All sequence data presented in this article is submitted and is available in the public repository of NCBI under Bioproject number PRJNA673998, experiment Biosample ID and Accession number of MAGs are shown in Table 1.

## Funding

This work was supported by funds from the Department of Biotechnology (DBT), National Bureau of Agriculturally Important Microorganisms (NBAIM), University Grants Commission-Career Advancement Scheme and Department of Science and Technology-Purse grant. S.N. and M.B. thank Council of Scientific and Industrial Research (CSIR) for providing doctoral fellowships.

## Author’s contribution

SN and CT planned the study. SN performed the analysis. SN and CT wrote the manuscript. MM performed the physicochemical analysis, HR performed the Scanning Electron Microscopy, MS, RL and RKN critically reviewed the manuscript and improved it. All authors read and approved the final manuscript.

## Conflict of Interest

The authors declare that they have no conflict of interest.

## Supplementary material legends

**Supplementary File S1**: Accession number of closest neighbors of MAGs available in NCBI. Functional COG subsystem distribution in 41 MAGs.

**Supplementary File S2**: The representation of distribution of gene involved in alkylphosphonate utilization, denitrification, denitrifying gene clusters, nitrosative stress and sulfate reductase complex.

**Supplementary File S3:** The gene names involved in Nitrogen, Phosphorous, sulfur and iron metabolism.

**Supplementary File S4:** The gene names involved in metal resistance and tolerance such as Chromium, Cadmium, Zinc, Cobalt, Copper and Arsenic.

**Supplementary File S5:** The Biogeochemical cycling entropy scores distribution in MAGs and reconstructed percentage of metabolic pathways involved in biogeochemical cycling.

## References

1. Agency for Toxic Substances and Disease Registry (ATSDR) Toxicological Profile for Copper. Atlanta, GA: Centers for Disease Control; 2002.

2. Ahemad M., (2012) Implications of bacterial resistance against heavy metals in bioremediation: A review. IIOABJ 3, 39–46

3. Alcamán-Arias, M. E. et al. Diurnal changes in active carbon and nitrogen pathways along the temperature gradient in porcelana hot spring microbial mat. Front. Microbiol. 9, 2353 (2018).

4. Altschul, S.F., Gish, W., Miller, W., Myers, E.W., Lipman, D.J. (1990) Basic local alignment search tool. J Mol Biol 215:403–410.

5. Anantharaman, K., Hausmann, B., Jungbluth, S.P. et al. Expanded diversity of microbial groups that shape the dissimilatory sulfur cycle. ISME J 12, 1715–1728 (2018). https://doi.org/10.1038/s41396-018-0078-0

6. Andreini C, Banci L, Bertini I, Rosato A. (2008) Occurrence of copper proteins through the three domains of life: a bioinformatic approach. J Proteome Res 7:209–216.

7. Bhattacharjee H, Zhou T, Li J, Gatti DL, Walmsley AR, Rosen BP. Structure-function relationships in an anion-translocating ATPase. Biochem Soc Trans. 2000;28(4):520–6.

8. Bowers, R., Kyrpides, N., Stepanauskas, R. et al. Minimum information about a single amplified genome (MISAG) and a metagenome-assembled genome (MIMAG) of bacteria and archaea. Nat Biotechnol 35, 725–731 (2017). https://doi.org/10.1038/nbt.3893

9. Brettin, T., Davis, J., Disz, T. et al. RASTtk: A modular and extensible implementation of the RAST algorithm for building custom annotation pipelines and annotating batches of genomes. Sci Rep 5, 8365 (2015). https://doi.org/10.1038/srep08365

10. Caetano, T., van der Donk, W., & Mendo, S. (2020). Bacteroidetes can be a rich source of novel lanthipeptides: The case study of Pedobacter lusitanus. Microbiological research, 235, 126441.

11. Chaumeil PA, et al. 2019. GTDB-Tk: A toolkit to classify genomes with the Genome Taxonomy Database. Bioinformatics, btz848.

12. Cowan, D., Tuffin, M., Mulako, I., & Cass, J. (2012). 12 Terrestrial Hydrothermal Environments.

13. De Anda V, Zapata-Peñasco I, Poot-Hernandez AC, Eguiarte LE, Contreras-Moreira B, Souza V. MEBS, a software platform to evaluate large (meta)genomic collections according to their metabolic machinery: unraveling the sulfur cycle. Gigascience. 2017 Nov 1;6(11):1–17. doi: 10.1093/gigascience/gix096.

14. Dodsworth, J. A., Hungate, B. A. & Hedlund, B. P. Ammonia oxidation, denitrification and dissimilatory nitrate reduction to ammonium in two US Great Basin hot springs with abundant ammonia-oxidizing archaea. Environ. Microbiol. 13, 2371–2386 (2011).

15. Dunbar, W. S. (2017). Biotechnology and the mine of tomorrow. Trends Biotechnol. 35, 79–89. doi: 10.1016/j.tibtech.2016.07.004

16. Franke, S., Grass, G., Rensing, C., Nies, D.H., 2003. Molecular analysis of the coppertransporting efflux system CusCFBA of Escherichia coli. J. Bacteriol. 185, 3804–3812.

17. Giacomucci, L., Purdy, K. J., Zanardini, E., Polo, A. & Cappitelli, F. A new non-degenerate primer pair for the specific detection of the nitrite reductase gene nrfA in the genus *Desulfovibrio*. J. Mol. Microbiol. Biotechnol. 22, 345–351 (2012).

18. Gupta, N., Balomajumder, C., & Agarwal, V. K. (2010). Enzymatic mechanism and biochemistry for cyanide degradation: a review. Journal of hazardous materials, 176(1-3), 1–13.

19. Harvey LJ, McArdle HJ. Biomarkers of copper status: a brief update. Br J Nutr. 2008; 99(S3): S10–S13.

20. Hedlund, B.P., Gosink, J.J. & Staley, J.T. Verrucomicrobia div. nov., a new division of the Bacteria containing three new species of Prosthecobacter. Antonie Van Leeuwenhoek 72, 29–38 (1997). https://doi.org/10.1023/A:1000348616863

21. Heng Li, Bob Handsaker, Alec Wysoker, Tim Fennell, Jue Ruan, Nils Homer, Gabor Marth, Goncalo Abecasis, Richard Durbin, 1000 Genome Project Data Processing Subgroup, The Sequence Alignment/Map format and SAMtools, Bioinformatics, Volume 25, Issue 16, 15 August 2009, Pages 2078–2079, https://doi.org/10.1093/bioinformatics/btp352

22. Hibbing ME, Fuqua C, Parsek MR, Peterson SB. 2010. Bacterial competition: surviving and thriving in the microbial jungle. Nat Rev Microbiol 8:15–25.

23. Hirota R, Kuroda A, Kato J, Ohtake H. Bacterial phosphate metabolism and its application to phosphorus recovery and industrial bioprocesses. J Biosci Bioeng. 2010 May;109(5):423–32. doi: 10.1016/j.jbiosc.2009.10.018.

24. Hua, ZS., Wang, YL., Evans, P.N. et al. (2019) Insights into the ecological roles and evolution of methyl-coenzyme M reductase-containing hot spring Archaea. Nat Commun 10, 4574. https://doi.org/10.1038/s41467-019-12574-y

25. Hyatt D, Chen GL, Locascio PF, Land ML, Larimer FW, Hauser LJ. Prodigal: prokaryotic gene recognition and translation initiation site identification. BMC Bioinformatics. 2010 Mar 8;11:119. doi: 10.1186/1471-2105-11-119.

26. Jacoby RP, Succurro A and Kopriva S (2020) Nitrogen Substrate Utilization in Three Rhizosphere Bacterial Strains Investigated Using Proteomics. Front. Microbiol. 11:784. doi: 10.3389/fmicb.2020.00784

27. Jain, C., Rodriguez-R, L.M., Phillippy, A.M., Konstantinidis K.T. and Aluru S. High throughput ANI analysis of 90K prokaryotic genomes reveals clear species boundaries. Nat Commun 9, 5114 (2018). https://doi.org/10.1038/s41467-018-07641-9

28. Jochum LM, Schreiber L, Marshall IPG, Jørgensen BB, Schramm A, Kjeldsen KU (2018) Single-cell genomics reveals a diverse metabolic potential of uncultivated Desulfatiglans-related Deltaproteobacteria widely distributed in marine sediment. Front Microbiol. 9:2038.

29. Kang D, Li F, Kirton ES, Thomas A, Egan RS, An H, Wang Z. 2019. MetaBAT 2: an adaptive binning algorithm for robust and efficient genome reconstruction from metagenome assemblies. PeerJ Preprints 7:e27522v1

30. Kim BE, Nevitt T, Thiele DJ. 2008. Mechanisms for copper acquisition, distribution and regulation. Nat Chem Biol 4:176–185.

31. Kraft, B., Strous, M. & Tegetmeyer, H. E. Microbial nitrate respiration—Genes, enzymes and environmental distribution. J. Biotechnol. 155, 104–117 (2011).

32. Kunin V, Copeland A, Lapidus A, et al. (2008) A Bioinformatician’s guide to metagenomics. Microbiol Mol Biol Rev 72:557–78.

33. Kushkevych, I., Cejnar, J., Treml, J., Dordević, D., Kollar, P., & Vítězová, M. (2020). Recent Advances in Metabolic Pathways of Sulfate Reduction in Intestinal Bacteria. Cells, 9(3), 698. https://doi.org/10.3390/cells9030698

34. Lam, Phyllis and Kuypers, Marcel M. M. (2011). “Microbial Nitrogen Processes in Oxygen Minimum Zones”. Annual Review of Marine Science. 3: 317–345. Bibcode: 2011ARMS.…3..317L. doi:10.1146/annurev-marine-120709-142814.

35. Langmead, B., & Salzberg, S. L. (2012). Fast gapped-read alignment with Bowtie 2. Nature methods, 9(4), 357–359. https://doi.org/10.1038/nmeth.1923

36. Li, SJ., Hua, ZS., Huang, LN. et al. Microbial communities evolve faster in extreme environments. Sci Rep 4, 6205 (2014). https://doi.org/10.1038/srep06205

37. Lin Y, Wang L, Xu K, Li K, Ren H. Revealing taxon-specific heavy metal-resistance mechanisms in denitrifying phosphorus removal sludge using genome-centric metaproteomics. Microbiome. 2021 Mar 22;9(1):67. doi: 10.1186/s40168-021-01016-x.

38. Lu, J., Zhu, B., Struewing, I. et al., (2019) Nitrogen-phosphorous-associated metabolic activities during the development of a cyanobacterial bloom revealed by metatranscriptomics. Sci Rep. 9, 2480. doi.org/10.1038/s41598-019-38481-2

39. Mark, P. (2018). Staphylococcal bacteriocins. In Pet-To-Man Travelling Staphylococci (pp. 161–171). Academic Press.

40. Mathur, J., Bizzoco, R. W., Ellis, D. G., Lipson, D. A., Poole, A. W., Levine, R., & Kelley, S. T. (2007). Effects of abiotic factors on the phylogenetic diversity of bacterial communities in acidic thermal springs. Applied and environmental microbiology, 73(8), 2612–2623.

41. Matsen, F.A., Kodner, R.B. & Armbrust, E. pplacer: linear time maximum-likelihood and Bayesian phylogenetic placement of sequences onto a fixed reference tree. BMC Bioinformatics 11, 538 (2010). https://doi.org/10.1186/1471-2105-11-538

42. McDaniel EA, Wever R, Oyserman BO, Noguera DR, McMahon KD. Genome-Resolved Metagenomics of a Photosynthetic Bioreactor Performing Biological Nutrient Removal. Microbiol Resour Announc. 2021 May 6;10(18):e00244–21. doi: 10.1128/MRA.00244-21.

43. Merino N, Aronson HS, Bojanova DP, Feyhl-Buska J, Wong ML, Zhang S and Giovannelli D (2019) Living at the Extremes: Extremophiles and the Limits of Life in a Planetary Context. Front. Microbiol. 10:780. doi: 10.3389/fmicb.2019.00780.

44. Micallef M. L., D’Agostino P. M., Sharma D., Viswanathan R., Moffitt M. C. (2015). Genome mining for natural product biosynthetic gene clusters in the Subsection V cyanobacteria. BMC Genomics 16:669 10.1186/s12864-015-1855-z.

45. Mohan, S. B., Schmid, M., Jetten, M. & Cole, J. Detection and widespread distribution of the nrfA gene encoding nitrite reduction to ammonia, a short circuit in the biological nitrogen cycle that competes with denitrification. FEMS Microbiol. Ecol. 49, 433–443 (2004).

46. Nagar S, Talwar C, Motelica-Heino M, Richnow HH, Shakarad M, Lal R, Negi RK (2021a) Microbial ecology of sulfur biogeochemical cycling at a mesothermic hot spring atop Northern Himalayas, India. bioRxiv. Preprint. https://doi.org/10.1101/2021.12.02.470874

47. Nagar S, Talwar C, Bharti M, Yadav S, Siwach S, Negi RK (2021b) Metagenome-assembled genomes recovered from the datasets of a high-altitude Himalayan hot spring Khirganga, Himachal Pradesh, India. Data in Brief. 39:107551.

48. Nagatani, H., Shimizu, M. & Valentine, R. C. The mechanism of ammonia assimilation in nitrogen fixing bacteria. Arch. Mikrobiol. 79, 164–175 (1971).

49. Nanda M, Kumar V, Sharma DK. Multimetal tolerance mechanisms in bacteria: The resistance strategies acquired by bacteria that can be exploited to ‘clean-up’ heavy metal contaminants from water. Aquat Toxicol. 2019 Jul; 212:1–10. doi: 10.1016/j.aquatox.2019.04.011.

50. Nasreen Amin, Vinod K. Kannaujiya, in Evolutionary Diversity as a Source for Anticancer Molecules, 2021

51. Norton, J. M., Alzerreca, J. J., Suwa, Y., & Klotz, M. G. (2002). Diversity of ammonia monooxygenase operon in autotrophic ammonia-oxidizing bacteria. Archives of microbiology, 177(2), 139–149.

52. Norton, J. M., Klotz, M. G., Stein, L. Y., Arp, D. J., Bottomley, P. J., Chain, P. S.,… & Starkenburg, S. R. (2008). Complete genome sequence of Nitrosospira multiformis, an ammonia-oxidizing bacterium from the soil environment. Applied and environmental microbiology, 74(11), 3559–3572.

53. O’Brien, J., and Wright, G. D. (2011) An ecological perspective of microbial secondary metabolism. Curr. Opin. Biotechnol. 22, 552–558. doi: 10.1016/j.copbio.2011.03.010

54. Outten, W., Huffman, D.L., Hale, J.A., O’Halloran, T.V., 2001. The independent cue and cus systems confer copper tolerance during aerobic and anaerobic growth in Escherichia coli. J. Biol. Chem. 276, 30670–30677.

55. Pal C, Bengtsson-Palme J, Rensing C, Kristiansson E, Larsson DG. BacMet: antibacterial biocide and metal resistance genes database. Nucleic Acids Res. 2014 Jan;42(Database issue):D737–43. doi: 10.1093/nar/gkt1252.

56. Parks DH, Imelfort M, Skennerton CT, Hugenholtz P, Tyson GW. 2014. Assessing the quality of microbial genomes recovered from isolates, single cells, and metagenomes. Genome Research, 25: 1043–1055.

57. Parks, D.H., et al. 2020. A complete domain-to-species taxonomy for Bacteria and Archaea. Nature Biotechnology, https://doi.org/10.1038/s41587-020-0501-8.

58. Pearson, A., Pi, Y., Zhao, W., Li, W., Li, Y., Inskeep, W.,… & Zhang, C. L. (2008). Factors controlling the distribution of archaeal tetraethers in terrestrial hot springs. Applied and Environmental Microbiology, 74(11), 3523–3532.

59. Pettit, R. K. (2011). Small-molecule elicitation of microbial secondary metabolites. Microb. Biotechnol. 4, 471–478. doi: 10.1111/j.1751-7915.2010.00196.x

60. Ridley, C. P., Lee, H. Y., & Khosla, C. (2008). Evolution of polyketide synthases in bacteria. Proceedings of the National Academy of Sciences, 105(12), 4595–4600.

61. Rohwerder, T. (2020). New structural insights into bacterial sulfoacetaldehyde and taurine metabolism. Biochemical Journal, 477(8), 1367–1371.

62. Ruiz, B., Chávez, A., Forero, A., García-Huante, Y., Romero, A., Sánchez, M., et al. (2010) Production of microbial secondary metabolites: regulation by the carbon source. Crit. Rev. Microbiol. 36, 146–167. doi: 10.3109/10408410903489576.

63. Seemann T.: rapid prokaryotic genome annotation Bioinformatics 2014 Jul 15;30(14):2068–9. PMID:24642063

64. Segata N, Börnigen D, Morgan XC, Huttenhower C. PhyloPhlAn is a new method for improved phylogenetic and taxonomic placement of microbes. Nat Commun. 2013;4:2304.

65. Sheik CS, Jain S, Dick GJ. Metabolic flexibility of enigmatic SAR324 revealed through metagenomics and metatranscriptomics. Environ Microbiol. 2014;16:304–17.

66. Skirnisdottir, S., Hreggvidsson, G. O., Hjörleifsdottir, S., Marteinsson, V. T., Petursdottir, S. K., Holst, O., & Kristjansson, J. K. (2000). Influence of sulfide and temperature on species composition and community structure of hot spring microbial mats. Applied and Environmental Microbiology, 66(7), 2835–2841.

67. Song, S., Jia, Z., Xu, J., Zhang, Z., & Bian, Z. (2011). N-butyryl-homoserine lactone, a bacterial quorum-sensing signaling molecule, induces intracellular calcium elevation in Arabidopsis root cells. Biochemical and biophysical research communications, 414(2), 355–360.

68. Srivastava S, Dong H, Briggs BR (2020) The Effect of Spring Water Geochemistry on Copper Proteins in Tengchong Hot Springs, China. Appl Environ Microbiol.; 86(13):e00581–20. doi: 10.1128/AEM.00581-20.

69. Stern BR. Essentiality and toxicity in copper health risk assessment: overview, update and regulatory considerations. Toxicol Environ Health A. 2010;73(2):114–127.

70. Tripathy, S., Padhi, S. K., Mohanty, S., Samanta, M. & Maiti, N. K. Analysis of the metatranscriptome of microbial communities of an alkaline hot sulfur spring revealed different gene encoding pathway enzymes associated with energy metabolism. Extremophiles 20, 525–536 (2016).

71. Van Thuoc, D., Hien, T. T., & Sudesh, K. (2019). Identification and characterization of ectoine-producing bacteria isolated from Can Gio mangrove soil in Vietnam. Annals of Microbiology, 69(8), 819–828.

72. Wanner, B. L.: Molecular genetics of carbon-phosphorus bond cleavage in bacteria, Biodegradation, 5, 175–184 (1994).

73. Weinitschke, S. (2010). New intermediates, pathways, enzymes and genes in the microbial metabolism of organosulfonates (Doctoral dissertation).

74. White AK, Metcalf WW. Microbial metabolism of reduced phosphorus compounds. Annu Rev Microbiol. 2007; 61:379–400. doi: 10.1146/annurev.micro.61.080706.093357.

75. White AK, Metcalf WW. 2004. The *htx* and *ptx* operons of *Pseudomonas stutzeri* WM88 are new members of the Pho regulon. J. Bacteriol. 186:587

76. Wilkins, L.G.E., Ettinger, C.L., Jospin, G. et al. Metagenome-assembled genomes provide new insight into the microbial diversity of two thermal pools in Kamchatka, Russia. Sci Rep 9, 3059 (2019). https://doi.org/10.1038/s41598-019-39576-6

77. Willsky GR, Malamy MH. 1976. Control of the synthesis of alkaline phosphatase and the phosphate-binding protein in *Escherichia coli*. J. Bacteriol. 127:595–609

78. Xu Z, Zhou G. [Research advance in nitrogen metabolism of plant and its environmental regulation]. Ying Yong Sheng Tai Xue Bao. 2004 Mar;15(3):511–6. Chinese. PMID: 15228008.

79. Xu, F., Wu, Y., Zhang, C., Davis, K. M., Moon, K., Bushin, L. B., et al. (2019) A genetics-free method for high-throughput discovery of cryptic microbial metabolites. Nat. Chem. Biol. 15, 161–168. doi: 10.1038/s41589-018-0193-2.

80. Yeoh YK, Sekiguchi Y, Parks DH, et al. (2015) Comparative genomics of candidate phylum TM6 suggests that parasitism is widespread and ancestral in this lineage. Mol Biol Evol 33:915–27.

81. Yilmaz P, Yarza P, Rapp JZ, Glöckner FO. (2016) Expanding the world of marine bacterial and archaeal clades. Front Microbiol.; 6:1524.

82. Zhang, M., and Wang, H. (2016). Preparation of immobilized sulfate reducing bacteria (SRB) granules for effective bioremediation of acid mine drainage and bacterial community analysis. Minerals Eng. 92, 63–71. doi: 10.1016/j.mineng.2016.02.008.

83. Zheng L, Ren M, Xie E, Ding A, Liu Y, Deng S and Zhang D (2019) Roles of Phosphorus Sources in Microbial Community Assembly for the Removal of Organic Matters and Ammonia in Activated Sludge. Front. Microbiol. 10:1023. doi: 10.3389/fmicb.2019.01023.

